# Learning-induced plasticity in the barrel cortex is disrupted by inhibition of layer 4 somatostatin-containing interneurons

**DOI:** 10.1101/2020.05.11.087791

**Authors:** G Dobrzanski, A Lukomska, R Zakrzewska, A Posluszny, D Kanigowski, J Urban-Ciecko, M Liguz-Lecznar, M Kossut

## Abstract

Learning-related plasticity in the cerebral cortex is linked to the action of disinhibitory circuits of interneurons. Pavlovian conditioning, in which stimulation of the vibrissae is used as conditioned stimulus, induces plastic enlargement of the cortical functional representation of vibrissae activated during conditioning, visualized with [^14^C]-2-deoxyglucose (2DG). Using layer-specific, cell-selective DREADD transductions, we examined the involvement of somatostatin- (SOM-INs) and vasoactive intestinal peptide (VIP-INs)-containing interneurons in the development of learning-related plastic changes. We injected DREADD-expressing vectors into layer IV (L4) barrels or layer II/III (L2/3) areas corresponding to activated vibrissae. The activity of interneurons was modulated during training, and 2DG maps were obtained 24 h later. In mice with L4 but not L2/3 SOM-INs suppressed during conditioning, the plastic change of whisker representation and the conditioned reaction were absent. No effect of inhibiting VIP-INs was found. We report that the activity of L4 SOM-INs is indispensable for learning-induced plastic change.

## INTRODUCTION

Learning, even in a simple conditioning paradigm, modifies the representation of encoded stimuli (Blake et al., 2006; Froemke et al., 2013, 2007; Gilbert et al., 2009; Kilgard and Merzenich, 2002; Rosselet et al., 2011; Weinberger and Bakin, 1998). We have described a learning-dependent plastic change of the functional representation of vibrissae in the mouse barrel cortex following conditioning consisting of pairing stimulation of a row of vibrissae with tail shock (Siucinska and Kossut, 1996). Conditioning resulted in an enlargement of the cortical representation of the cognate row of whiskers, as visualized with [^14^C]-2-deoxyglucose autoradiography (2DG), mainly in cortical layer IV (L4). This plastic change was associated with upregulation of the glutamic acid decarboxylase level in interneurons containing somatostatin (SOM-INs) in L4 of the “trained” whisker representation (Cybulska-Klosowicz et al., 2013b) and with an increased number of inhibitory synapses on double synapse spines in the same location (Jasinska et al., 2016), likely made by SOM-INs (Chiu et al., 2013).

SOM-INs consist of several classes of cells with diverse physiologies, morphologies, and synaptic targets. SOM-INs from cortical layers 2/3/5 and L4 SOM-INs represent different morphological types (Martinotti *vs.* non-Martinotti, respectively) and have complementary patterns of synaptic input and axonal output, which suggests that they are differentially engaged by neuronal activity and selectively control specific layers (Muñoz et al., 2017; Naka et al., 2019). SOM-INs regulate the activity of cortical projection neurons either by direct inhibition or by disinhibition (Cottam et al., 2013; Karnani et al., 2014; Letzkus et al., 2011). Generally, Martinotti SOM-INs target apical dendrites of pyramidal neurons, regulating their activities by controlling dendritic integration of synaptic inputs (Berger et al., 2010; Gentet et al., 2012). L4 SOM-INs target projection neurons of L4 and fast-spiking parvalbumin interneurons (PV-INs) to release thalamorecipient excitatory projection neurons from inhibition and enhance information flow into the upper cortical layers (Xu et al., 2013). The mechanism of disinhibition is particularly important during learning and plasticity, and the role of SOM-INs in these phenomena has been intensively investigated (Wolff et al., 2014; Xu et al., 2019). In the visual cortex, visual discrimination reduced the activity of L3 SOM-INs (Makino and Komiyama, 2015) but enhanced their stimulus selectivity, implicating them in gating selectivity changes (Khan et al., 2018). The activity of SOM-INs is strongly modulated by acetylcholine (N. Chen et al., 2015; Urban-Ciecko et al., 2018) and by inhibitory input from vasoactive intestinal polypeptide-containing interneurons (Pfeffer et al., 2013). The L2/3 VIP-SOM circuit was shown to be a potent regulator of cortical activity and learning (Fu et al., 2015; Lee et al., 2013; Pi et al., 2013). Due to disinhibitory cortical circuits, “blanket inhibition” extended by SOM-INs can be locally and momentarily curtailed by the action of VIP-INs, thus facilitating learning-related synaptic plasticity in projection neurons (Karnani et al., 2016).

The role of L4 SOM-INs in cortical plasticity has not been extensively examined thus far. Here, we investigated how selective, layer-specific, chemogenetic modulation of SOM-IN activity shapes learning-induced plasticity in the barrel cortex of mice and whether silencing L2/3 VIP-INs can modulate the formation of this plastic change.

## METHODS

### Subjects

Transgenic SOM-IRES-Cre, VIP-IRES-Cre (Taniguchi et al., 2011) and Ai14 (Madisen et al., 2010) mouse lines (stock: 013044, 010908 and 007914, respectively) were acquired from The Jackson Laboratory (USA) and bred in the Animal Facility of the Nencki Institute of Experimental Biology PAS (Warsaw, Poland). All mice used in the experiments were male 2-to 3-month-old heterozygotes. Genotyping of SOM-Cre and VIP-Cre mice was performed according to protocols provided by The Jackson Laboratory. During validation experiments, homozygous SOM-Cre and VIP-Cre mice were crossed with the Ai14 line to obtain tdTomato expression following Cre-mediated recombination. Before the experiments, all animals were housed in cages with nesting material, four-five per cage, in a temperature- and humidity-controlled room (20–22 °C, 40–50% humidity), on a 12 h light/dark cycle (lights on at 7 a.m.), with *ad libitum* access to food and water. After surgery, the mice were housed separately under the same conditions. All procedures were consistent with the European Community Council Directive (2010/63/UE) and were approved by the first Local Ethical Committee in Warsaw (456/2017 and 279/2017). All efforts were made to minimize the number of animals used and their suffering.

### Optical imaging recordings

Intrinsic signals from the barrel cortex were imaged using a pair of front-to-front camera lenses (Nikkor 50 mm f/1.2, Nikon) on a complementary metal oxide semiconductor (CMOS) camera (Photon Focus MV1-D1312-160-CL-12) with a maximal resolution of 1312 x 1082 pixels and a pixel size of 8 x 8 μm. Synchronization of image acquisition with whisker stimulation was controlled with the imaging system Imager 3001 (Optical Imaging, Rehovot, Israel).

Mice were placed in a plexiglass box and initially anesthetized by inhalation of 3% isoflurane (Aerrane, Baxter). During the surgery and the recording, ~2% and ~1.5% isoflurane was provided, respectively. Body temperature was maintained at 37 °C (Harvard Apparatus, Cambridge, UK), and the breathing rate was monitored (Datex Capnomac Ultima, Finland). Before recording, the mice were subcutaneously injected with dexamethasone (0.2 mg/kg) and tolfenamic acid (4 mg/kg). Lidocaine solution (2%, 0.1 ml) was applied subcutaneously before exposing the skull. The skull was covered with a solution of agarose in saline (2.5%) to make the skull transparent and sealed with a coverslip. After recording, the agarose window was removed, and the mice received subcutaneous injections of tolfenamic acid (0.1 mg/mouse).

At the beginning of the recording, the skull over the barrel cortex was illuminated with 546 nm (green light) to capture the pattern of superficial blood vessels. Then, the camera focus was set at 400 μm beneath the pial surface, and functional imaging was performed with red light illumination (630 nm). Row B whiskers were activated to record their cortical representation. For a functional map, the vibrissa-barrel system was activated by displacing whiskers in the rostrocaudal direction with a frequency of 10 Hz for 6 s under the control of Master 8, a pulse stimulator triggered by the imaging system. Images of the pial surface with blood vessels and images of row B whisker activation were superimposed on each other, providing the visualization necessary for targeted viral injection.

### Viral injections

Under optical imaging, mice were injected with Cre-dependent serotype 2/2 adeno-associated viruses expressing inhibitory DREADDs, excitatory DREADDs or red fluorescent protein mCherry (4.9×10^12^ vg/ml of pssAAV-2/2-hSyn1-dlox-hM4D(Gi)_mCherry(rev)-dlox-WPRE-hGHp(A); 2.8×10^12^ vg/ml of pssAAV-2/2-hSyn1-dlox-hM3D(Gq)_mCherry(rev)-dlox-WPRE-hGHp(A) or 5.4×10^12^ vg/ml pssAAV-2/2-hSyn1-dlox-mCherry(rev)-dlox-WPRE-hGHp(A), respectively). Vectors were provided by the Viral Vector Facility, University of Zurich.

The skull above row B barrels was thinned with a dental drill, and a small fragment of the skull was lifted to produce a small entrance for the injection capillary. Row B barrels were injected with viral vectors (40 nl/barrel) using a nanoliter injector (Nanoliter 2010, WPI) ended with a glass capillary backfilled with paraffin oil (catalog: 76235, Sigma-Aldrich), one injection to each barrel. Cortical injections were performed perpendicularly to the surface of the brain at a depth of 150 (L2/3) or 330 μm (L4) from the brain surface, with a flow rate set at 4 nl/min. To prevent backflow of the viruses, after injection, we left the capillary in place for ten minutes and then slowly withdrew it. After the procedure, the skin was sutured with an absorbable suture (Dafilon, Braun, Germany), and the mice were subcutaneously injected with enrofloxacin (5 mg/kg) and tolfenamic acid (4 mg/kg). These drugs were administered for three consecutive days.

### Habituation, conditioning and DREADD activation

Three days after injection, the mice started a 3-week habituation to the restraining holder (10 minutes per day), a period allowing for DREADD expression. The restraining holder allowed the mouse to be kept in one place while permitting free head movements. Classical conditioning comprised three 10-min sessions, one session per day. One session consisted of 40 trials, and during each trial, three 3 s strokes of the row B whiskers on one side of the snout were applied manually (CS, conditioned stimulus) and coupled with a mild tail shock (0.5 mA, 0.5 ms; UCS unconditioned stimulus) applied at the end of the third CS stimulus. All conditioning sessions were video recorded. Transduced mice from the experimental group received an intraperitoneal injection of CNO (dissolved in sterile saline to a dose of 1 mg/kg; catalog: BML-NS105, Enzo Life Sciences) 30 min before each session of the conditioning. Transduced control group mice received saline instead of CNO in the same manner as the experimental group.

### Behavioral analysis

Behavioral verification of associative learning was performed as described previously (Cybulska-Klosowicz et al., 2009). Briefly, the development of a conditioned reaction (minifreezing) consisting of halting movements toward the stimulus during CS application was filmed. The percentage of head movement associated with CS per minute as well as per day was calculated.

### The 2-deoxyglucose procedure (2DG)

On a day after the last session of the conditioning, the mice were placed in the restraining holder, and all whiskers except those in row B on both sides of the snout were trimmed. Then, the mice were intramuscularly injected with 0.175 ml of [^14^]C 2-deoxyglucose (American Radiolabeled Chemicals, Inc., specific activity 53 mCi/mmol), and both row B whiskers were manually stimulated along the anteroposterior axis in both directions for 30 min, with a frequency of 2 Hz. Next, the mice were anesthetized with a lethal dose of barbiturate (Vetbutal, Biowet, Poland) and briefly perfused with 4% paraformaldehyde solution in PBS (pH=7.4, Santa Cruz Biotechnology). Their brains were removed, and the hemispheres were separated and flattened. The hemispheres were then cut into 30 μm thick serial sections tangential to the barrel cortex on a cryostat (−20 °C), and these were placed, together with [^14^C] radioactive standards, against Kodak X-ray mammography film for 1 week.

Autoradiograms were scanned with a calibrated densitometer (GS-900™, Bio-Rad), and the scans were pseudocolored using Image Lab™ Software and then analyzed in ImageJ software. A calibration curve was created based on the absolute gray levels of the [^14^]C standards. The signals on all analyzed autoradiograms were within the linear range of this curve. The criterion for labeling was that the intensity of 2DG uptake was at least 15% higher than in the surrounding cortex, as described in Siucinska and Kossut (1996). Labeling of row B representation fulfilling this criterion was thresholded, and the width of the labeling across the row covering thresholded pixels was measured in μm. The width of the row B representation was measured in four sections of L4, and these four values were averaged for each hemisphere. The identification of L4 was performed on counterstained sections from which the autoradiograms were obtained. For analysis of the viral injections, every fourth tangential section of the transduced hemisphere was mounted on microscope glass using VECTASHIELD^®^ Antifade Mounting Medium with DAPI (catalog: H-1200, Vector Laboratories) and analyzed using fluorescence microscopy. Animals were included in the analyses if at least two row B barrels (including B1) were transduced. Plasticity was defined as a change in representation width, calculated as a ratio between a representation width of conditioned row B to a control one in the contralateral hemisphere.

### Immunohistochemistry

For confirmation of the selectivity of transgene expression, homozygous SOM-Cre and VIP-Cre mice were crossed with the Ai14 line that expresses the red fluorescent protein tdTomato in Cre recombinase-expressing cells, and immunofluorescence staining for SOM, VIP and PV was performed on these brain sections.

Two-month-old F1 mice were transcardially perfused with 0.01 M phosphate-buffered saline (PBS) following 4% paraformaldehyde solution in PBS, pH=7.4 (PFA, Santa Cruz Biotechnology). After perfusion, the brains were removed and postfixed in 4% PFA for 24 h at 4 °C, cryoprotected in a series of sucrose solutions in PBS (10%, 20%, and 30%; 24 h each; 4 °C), cooled in n-heptane, placed on dry ice and stored at −80 °C. Thirty-micrometer sections were cut in a coronal plane using a cryostat (Leica CM1860 UV).

Free-floating sections were blocked for 60 min at room temperature (RT) in PBS containing 10% donkey normal serum (catalog: D9663, Sigma-Aldrich), 5% bovine serum albumin (catalog: A7906, Sigma-Aldrich) and 0.1% Triton-X (catalog: T8787, Sigma-Aldrich). Then, the sections were incubated overnight with appropriate primary antibodies in blocking buffer at 4 °C. The next day, the samples were washed in PBS and incubated with a secondary antibody for 2 h at RT. Then, they were washed again in PBS and mounted using VECTASHIELD^®^ Antifade Mounting Medium with DAPI (catalog: H-1200, Vector Laboratories). The immunofluorescence signal was analyzed using confocal or fluorescence microscopy (Zeiss Cell Observer SD Spinning Disk Confocal Microscope or Nikon Eclipse 80i, respectively). The images were post-processed using ImageJ software.

The primary antibodies used in a study were as follows: a rabbit anti-somatostatin polyclonal antibody (1:500, catalog: H-106, Santa Cruz), a mouse anti-parvalbumin polyclonal antibody (1:1000, catalog: P3088, Sigma Aldrich) and a rabbit anti-vasoactive intestinal peptide (1:500, catalog: 9535-0204, Bio-Rad). The secondary antibodies used in a study were as follows: donkey anti-rabbit IgG (H+L, Highly Cross-Adsorbed, Alexa Fluor 488; 1:500, catalog: A-21206, Thermo Fisher Scientific) and donkey anti-mouse IgG (H+L, Highly Cross-Adsorbed, Alexa Fluor 488; 1:500, catalog: A-21202, Thermo Fisher Scientific).

### Patch-clamp recordings

SOM-Cre mice (1.5 months old) with virus injections were anesthetized, and the brains were removed for brain slice preparation. Brain slices of 350 μm thickness were cut along a 45° plane toward the midline to obtain “across-rows” slices (Finnerty et al., 1999). Slices were recovered and maintained at 24 °C in regular artificial cerebrospinal fluid (ACSF) (in mM): 119 NaCl, 2.5 KCl, 2 MgSO_4_, 2 CaCl_2_, 1 NaH_2_PO_4_, 26.2 NaHCO_3_, 11 glucose equilibrated with 95/5% O_2_/CO_2_.

Neurons were classified as SOM+ according to the fluorescence and spiking features in response to 500 ms suprathreshold intracellular current injection. Somata of the fluorescently labeled neurons in the primary somatosensory cortex were targeted for whole-cell patch-clamp recording with borosilicate glass electrodes (resistance 4-8 MΩ) filled with the electrode internal solution, composed of (in mM) the following: 125 potassium gluconate, 2 KCl, 10 HEPES, 0.5 EGTA, 4 MgATP, and 0.3 NaGTP, at pH 7.25-7.35, 290 mOsm. To obtain spontaneous activity of SOM-INs, we performed recordings in modified ACSF composed of (in mM) the following: 119 NaCl, 3.5 KCl, 0.5 MgSO_4_, 1 CaCl_2_, 1 NaH_2_PO_4_, 26.2 NaHCO_3_, 11 glucose equilibrated with 95/5% O_2_/CO_2_ at room temperature. Electrophysiological data were acquired by a Multiclamp 700B and Axon Digidata 1550B (Molecular Devices) acquisition interface. The data were filtered at 3 kHz, digitized at 10 kHz, and collected by Clampex 10.6. Input resistance was analyzed online. CNO (dissolved in ACSF to a concentration of 10 μM) was bath applied for at least 10 min before data acquisition to assess the drug effects on the basic electrophysiological properties of SOM neurons (resting membrane potential and spontaneous activity).

### Statistical analyses

Statistical analyses were performed using GraphPad Prism5 software (Inc.). The results are presented as the mean±SEM, and p<0.05 was considered statistically significant. In patchclamp experiments, population data were analyzed using a two-tailed paired t-test. In 2DG experiments, the between-group comparison of row B labeling was performed using two-way ANOVA with Bonferroni post-tests, while comparison between hemispheres within one group was performed with a two-tailed paired t-test. A comparison of plasticity between two groups was performed using a two-tailed nonpaired t-test. In behavioral experiments, repeated-measures ANOVA with Tukey’s multiple comparison tests was used to compare minifreezing between sessions in one group of animals, whereas minifreezing in all three sessions between two experimental groups was analyzed using two-way ANOVA with Bonferroni post-tests.

## RESULTS

### Row B barrel transduction

Using DREADD-expressing viral vectors, we developed stable transduction of row B SOM-INs in L4 and L2/3 and VIP-INs in L2/3 in the respective groups of mice. In most cases, as shown in **Fig. 1A and B**, transduction covered three row-B barrels (B1-B3), but animals with two transduced barrels (B1-B2) were also included in the analyses. Transduced cells of L4 were localized in both hollows and walls. In a separate experiment, tangential L4 brain sections were immunostained for SOM, proving that more than 90% of the transduced cells were positive for somatostatin (data not shown). We used AAV serotype 2/2 to confine the transduction to one row and minimize the spread of the virus (Watakabe et al., 2015), but occasionally, some individual cells were also observed outside row B (as shown in **Fig. 1A**).

**Fig. 1.**
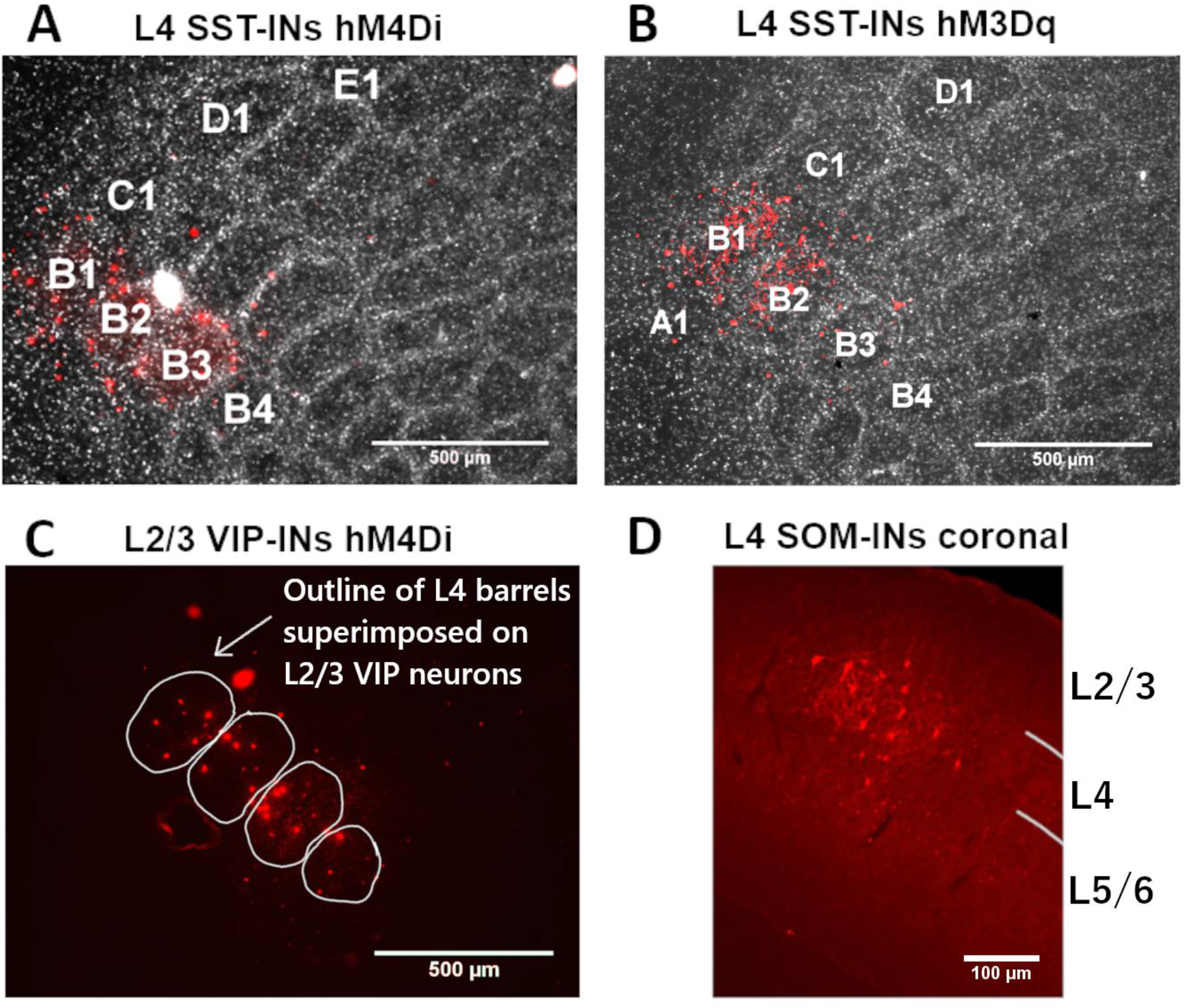
Transduction of row B barrels. **A**. Tangential section of the barrel cortex L4 showing transduced SOM-INs with inhibitory DREADDs (hM4Di-mCherry) in row B barrels colocalized with DAPI. Scale bar: 500 μm. **B.** Tangential section of the barrel cortex L4 showing transduced SOM-INs with excitatory DREADDs (hM3Dq-mCherry) in row B barrels colocalized with DAPI. Scale bar: 500 μm. **C.** Tangential section of the barrel cortex L2/3 showing transduced VIP-INs with inhibitory DREADDs (hM4Di-mCherry) in row B barrels. Localization of L2/3 VIP INs within row B columns was assessed by superimposing serial tangential sections of L4 and L2/3 aligned by blood vessels. Scale bar: 500 μm. **D.** Coronal section of the barrel cortex showing transduced SOM-INs with excitatory DREADDs (hM3Dq-mCherry) confined to L4. Scale bar: 100 μm.

Transduction was confined mostly to L4, as shown in the coronal section in **Fig. 1D**. We also analyzed transduction in other cortical layers and did not detect any fluorescent cells below L4 (in the L4 SOM-IN groups) or below L2/3 (in the VIP and L2/3 SOM-IN-transduced animals); sporadically, a few fluorescent cells along a capillary tract were detected in L2/3 (in the L4 SOM-IN-transduced animals). Localization of L2/3 VIP INs within row B columns was assessed by superimposing serial tangential sections of L4 and L2/3 aligned by blood vessels (**Fig. 1C)**.

### Control experiments and DREADD technique validation

#### Verification of transgene specificity

To verify whether Cre recombinase is cell-specific and to assess its ability to perform recombination *in vivo,* we crossed homozygous SOM-Cre and VIP-Cre mice with homozygous Ai14 mice to obtain tdTomato expression following Cre-mediated recombination. Coronal brain sections from either SOM-Ai14 or VIP-Ai14 mice were immunostained for all three main groups of cortical inhibitory interneurons (PV, SOM, VIP). The distribution across the brain (Neske et al., 2015) (Fig. 2). Qualitative analyses revealed colocalization between tdTomato and SOM or VIP (in the SOM-Ai14 and VIP-Ai14 mice, respectively), proving the transgene specificity. We did not observe any overlap between genetically labeled SOM or VIP cells and immunopositive cells within other types of interneurons, except for rarely observed, negligible in number parvalbumin immunosignals in the SOM-tdTomato cells, as has been reported before (Hu et al., 2013).

**Fig. 2.**
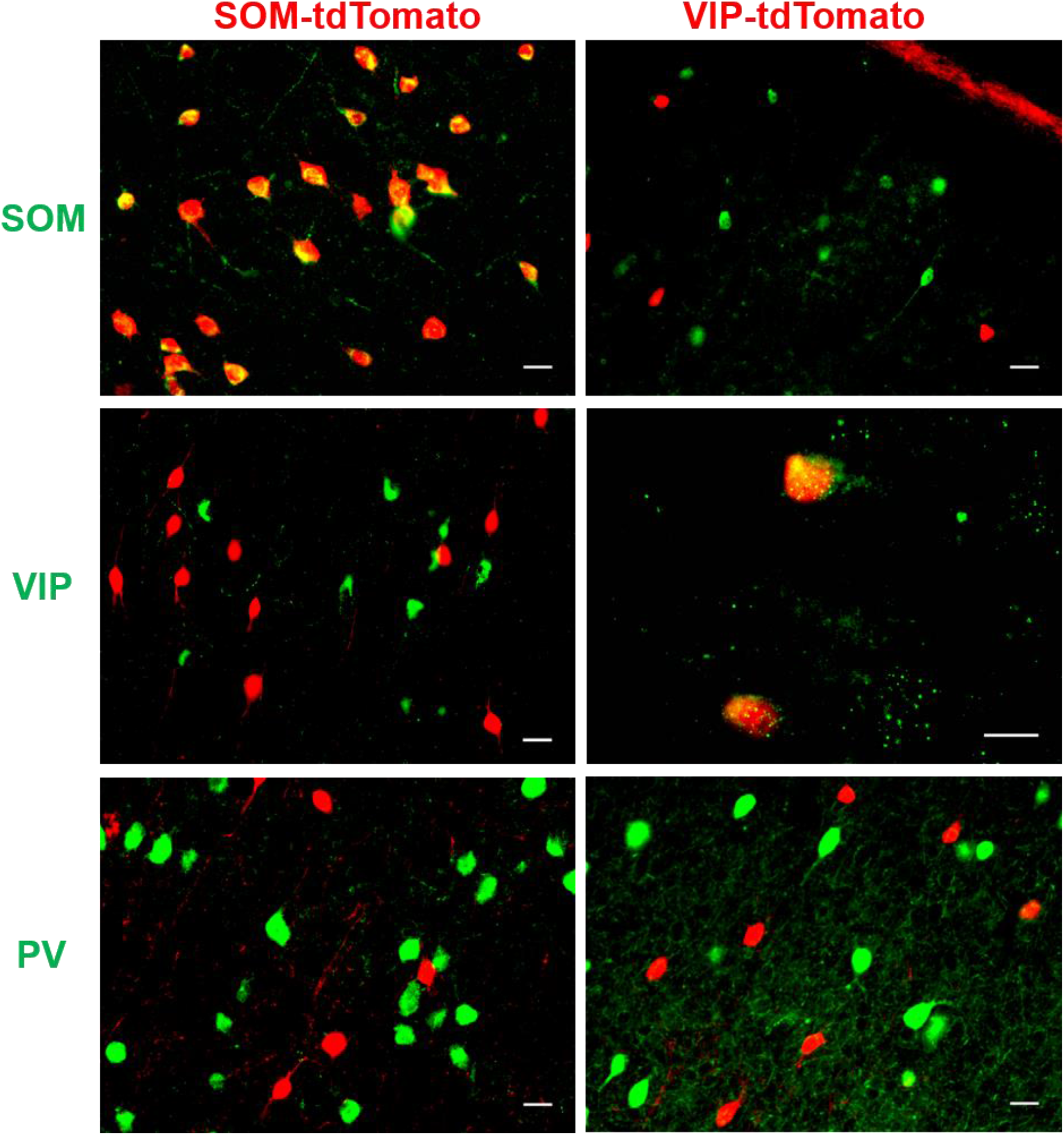
Immunofluorescent validation of transgene expression in SOM-Cre and VIP-Cre mice. Immunostaining for SOM, VIP, and PV in the mice expressing tdTomato in SOM or VIP INs. SOM and VIP immunostaining in the SOM-tdTomato and VIP-tdTomato mice, respectively, showed colocalization between tdTomato and fluorescent signals in the mouse barrel cortex. Immunostaining for VIP and PV (in SOM-tdTomato) and PV and SOM (in VIP-tdTomato mice) revealed no overlap between immunofluorescence from markers of particular types of interneurons and tdTomato. Scale bar: 10 μm.

#### Electrophysiological validation of DREADD activity *in vitro*

To validate the applicability of the DREADD technique in the modulation of interneuron activity, we performed whole-cell patch-clamp recordings of SOM-INs selectively transduced with DREADD-expressing viral vectors (hM4Di or hM3Dq) or empty vectors. Two weeks after injection, the mice were sacrificed, and the basic electrophysiological properties of the SOM-INs were recorded before and after CNO (10 μM) administration. This CNO concentration has been used *in vitro* to change the activity of neurons (Saloman et al., 2016), including GABAergic neurons (Koga et al., 2017). We found that CNO application did not influence the activity of the SOM-INs transduced with empty viruses but altered the activity of the cells transduced with DREADDs. Activation of hM4Di decreased the resting membrane potential (–61.74 ± 1.71 mV before and −65.82 ± 1.14 mV after CNO application, ** p=0.005, two-tailed paired t-test, n=8 cells) and the frequency of spontaneous firing of the SOM-INs (8.30 ± 4.28 Hz before and 2.55 ± 1.19 Hz after CNO application, *p=0.0313, two-tailed Wilcoxon test, n=8 cells). Activation of hM3Dq, however, provided an increase in both parameters (resting membrane potential: −55.08 ± 1.66 mV before and −52.08 ± 1.51 mV after CNO application, *** p<0.0001, two-tailed paired t-test, n=12 cells; frequency: 3.51 ± 1.12 Hz before and 6.49 ± 1.33 Hz after CNO application, ** p=0.0049, two-tailed Wilcoxon test, n=8 cells; (Fig. 3)). These *in vitro* results confirmed the regulatory effectiveness of the DREADD technique used in the experiments, and they indicate that the activity of interneurons within the mouse barrel cortex can be modulated with the chemogenetic DREADD approach in both directions.

**Fig. 3.**
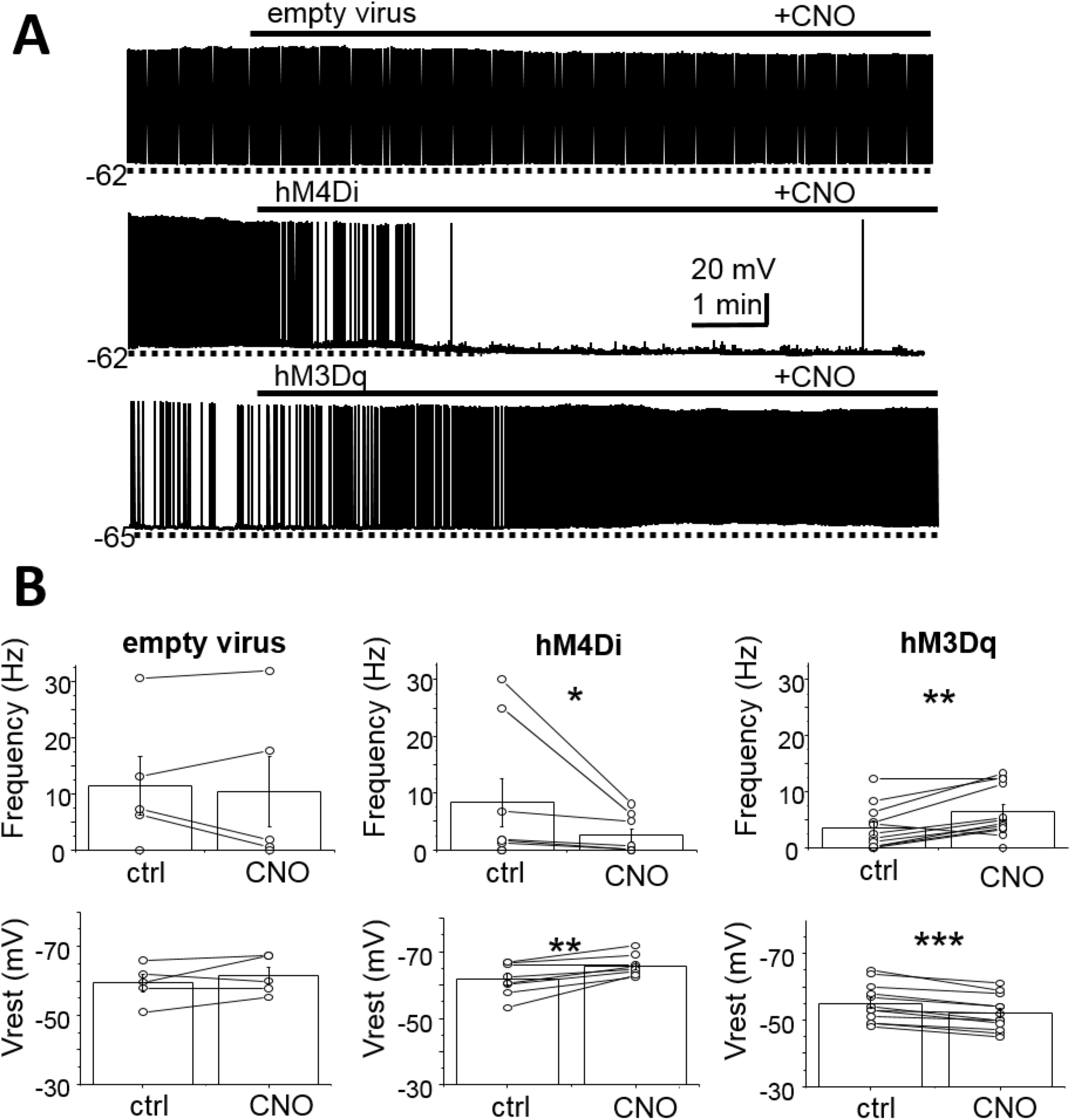
Electrophysiological validation of DREADD activity in SOM-INs *in vitro*. **A**. Examples of spontaneous activity of SOM interneurons transduced with empty viruses, inhibitory DREADDs (hM4Di), and excitatory DREADDs (hM3Dq), before and after CNO (10 μM) application. **B**. Mean frequency of spontaneous firing and resting membrane potential of SOM interneurons before and after CNO (10 μM) application in mice transduced with empty viruses or inhibitory or excitatory DREADDs. CNO did not change either the frequency (ns p=0.6295, two-tailed paired t-test, n=5 cells) or the resting potential (ns p=0.2677, two-tailed paired t-test, n=5 cells) in DREADD-free SOM-INs transduced with empty vectors. In the hM4Di-transduced SOM-INs, CNO application decreased the frequency (* p=0.0313, two-tailed Wilcoxon test, n=8 cells) as well as the resting potential (** p=0.005, two-tailed paired t-test, n=8 cells). In the hM3Dq-transduced SOM-INs, CNO increased both the frequency (** p=0.0049, two-tailed Wilcoxon test, n=8 cells) and the resting membrane potential (*** p<0.0001, two-tailed paired t-test, n=12 cells).

#### CNO alone does not influence learning-induced plastic changes

CNO was believed to be a selective and potent DREADD agonist as well as a pharmacologically inert drug. Because a study by MacLaren et al. (2016) showed that CNO in DREADD-free rats can disturb behavioral outcomes, we determined whether one of the most widely used doses of CNO in mice, 1 mg/kg, could influence the formation of plastic changes induced by classical conditioning. Two groups of SOM-Cre mice underwent three days of whisker-based fear conditioning, and 30 min before each session, one group of mice was intraperitoneally administered CNO (1 mg/kg), while the other group received a saline injection. After the 2DG experiment and autoradiography procedure, we compared the width of row B labeling between the conditioned and control hemispheres in both groups. Two-way ANOVA revealed the main effect of hemisphere (F_(1,24)_=59.29, p<0.0001) but not group (F_(1,24)_=1.433, p=0.2429) or interaction (F_(1,24)_=0.003738, p=0.9518) (Fig. 4). Bonferroni post-tests showed a significant difference between both hemispheres in row B representation width of the group receiving saline (### p<0.001, n=7) as well as CNO (### p<0.001, n=7), proving that in both groups of animals, plasticity was induced with similar intensity. Plasticity in the animals receiving saline equaled 1.245 ± 0.03 and in those receiving CNO was 1.249 ± 0.02 and did not differ between the groups (ns, p=0.9174, two-tailed paired t-test). We did not observe any nonspecific labeling in the CNO-treated animals compared to the saline-injected mice. Collectively, these results suggest that CNO (1 mg/kg) administration before each session of conditioning does not interfere with plastic change formation and can be applied as a DREADD actuator in our model of whisker-based classical conditioning. Therefore, we decided not to include another control group of mice transduced with an empty vector that received CNO.

**Fig. 4.**
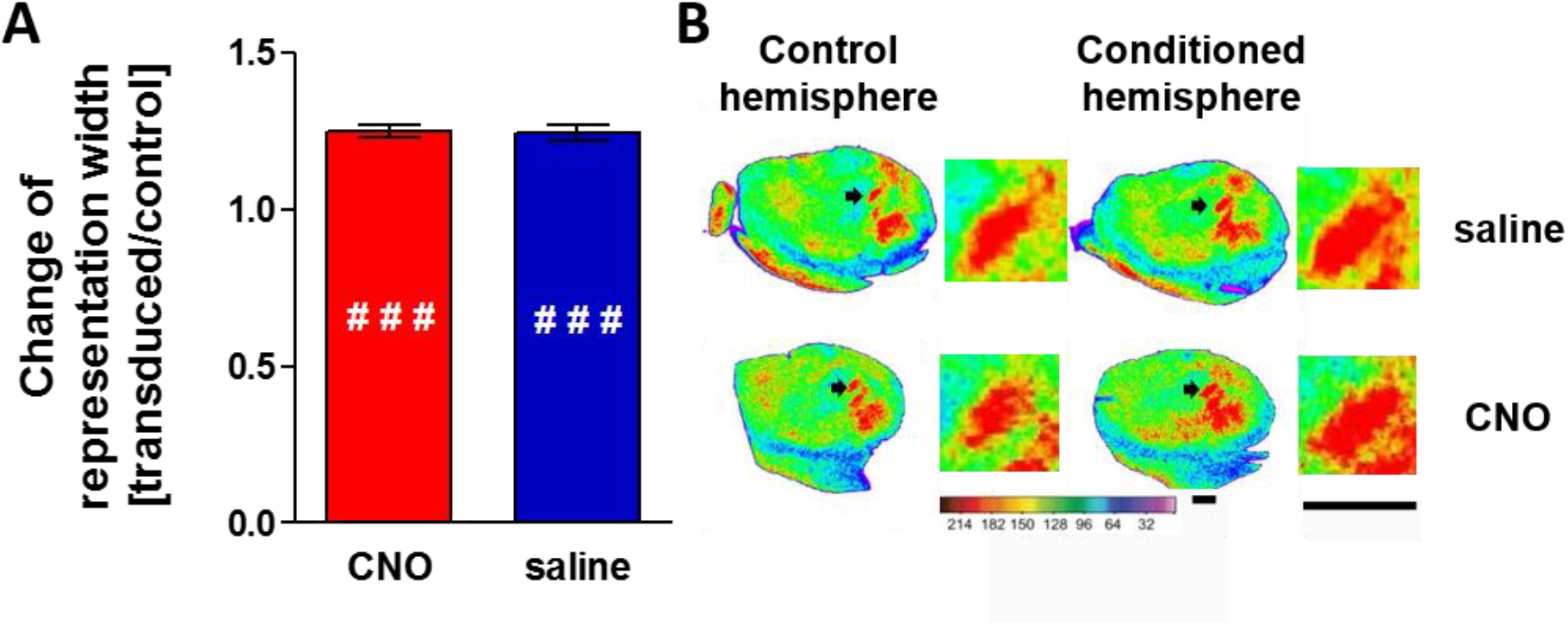
CNO alone does not influence learning-induced plastic changes. **A.** Plasticity shown as a change of representation width, calculated as a ratio between a representation width of conditioned row B to a control one in the opposite hemisphere (mean±SEM). Two-way ANOVA with Bonferroni post-tests showed a significant difference in row B width between the control and conditioned hemisphere (F(1,24)=59.29, p<0.0001) in the control group receiving saline (### p<0.001, n=7) as well in experimental group receiving CNO (### p<0.001, n=7). No significant difference in plasticity between the two groups (ns p=0.9174, two-tailed unpaired t-test) was observed. **B.** Examples of representative, pseudocolored autoradiograms (a pair of left control and right conditioned hemisphere shown in a row) taken from both the control (saline) and experimental groups (CNO). Black arrows indicate row B representations, magnified in a square on the right side. Scale bar = 1 mm.

### L4 SOM-IN inhibition blocks the formation of plastic changes induced by learning

To study how the blockade of SOM-IN activity during conditioning influences plasticity, we used the chemogenetic DREADD technique, which allows remote modulation of neuronal activity (Armbruster et al., 2007). The usefulness of the DREADD technique in changing SOM-IN activity *in vivo* was proven in 2014 (Soumier and Sibille, 2014), and since then, this approach has become widely used. Mice were injected with Cre-dependent vectors expressing hM4Di into the SOM-INs of L4 row B barrels. During conditioning, the experimental group received CNO (experimental group, hM4Di+CNO, n=6), while the control group received saline (control group, hM4Di+saline, n=6).

Analysis of the autoradiograms showed that in the CNO-treated hM4Di-transduced mice, the plastic change of trained row B representation was not induced. The row B representation width in the conditioned hemisphere was almost equal to the width in the control hemisphere, and this observation was consistent in all tested animals. In contrast, in the control group, conditioning-induced widening of the row B representation was observed. A significant difference in the width of row B representations was observed between the two groups (0.98 ± 0.2 and 1.23 ± 0.05, respectively; *** p<0.001, two-tailed unpaired t-test) (**Fig. 5A**).

**Fig. 5.**
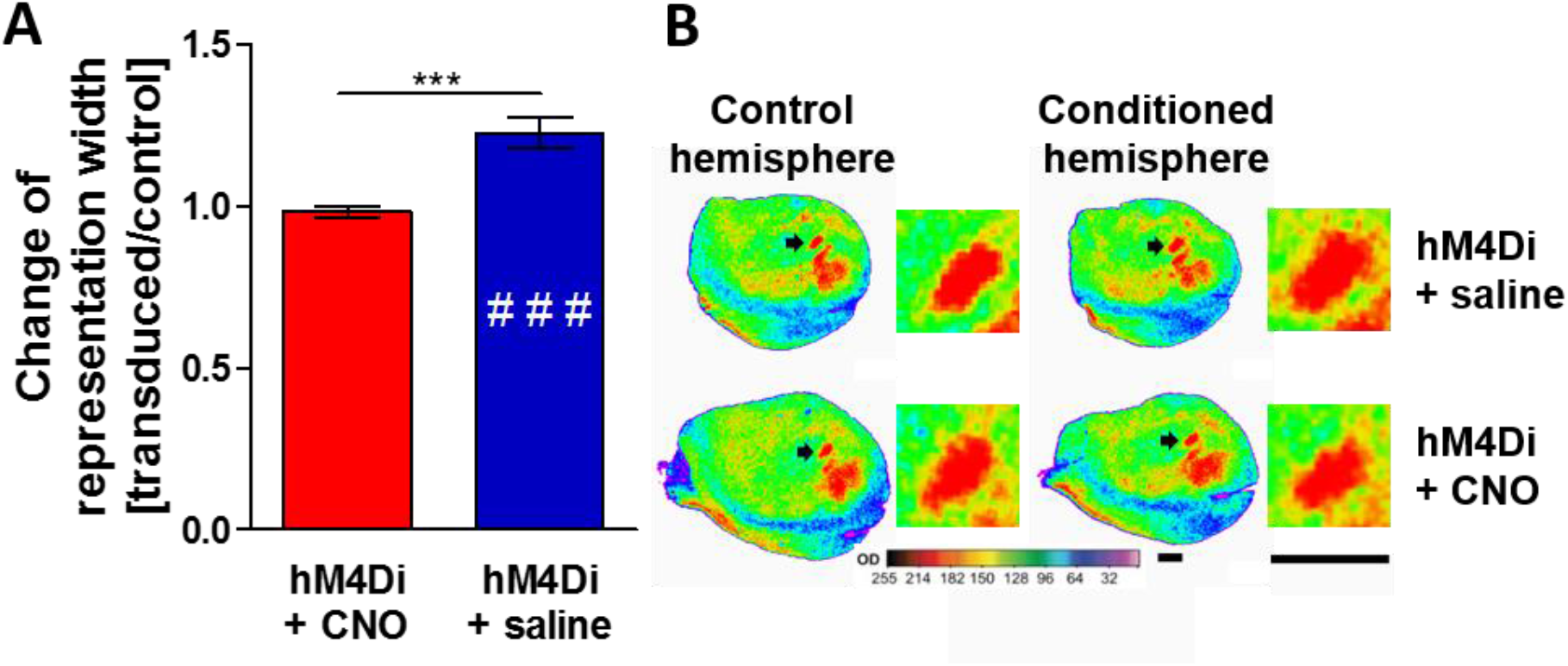
L4 SOM-IN inhibition blocks the formation of learning-induced plastic change. **A.** Plasticity shown as a change of representation width, calculated as a ratio between a representation width of conditioned (transduced) row B to a control one in the opposite hemisphere (mean±SEM). Two-way ANOVA with Bonferroni post-tests showed a significant difference in row B width between the control and conditioned hemisphere (F_(1,20)_=8.701, p=0.0079) in the control group (hM4Di+saline) (### p<0.001, n=6) but no change in SOM-IN inhibition (hM4Di+CNO) (ns p>0.05, n=6). A significant difference in plasticity between the two groups (*** p<0.0001, two-tailed unpaired t-test) was observed. **B.** Examples of representative, pseudocolored autoradiograms (a pair of left control and right conditioned hemisphere shown in a row) taken from both the control (hM4Di + saline) and experimental groups (hM4Di + CNO). Black arrows indicate row B representations, magnified in a square on the right side. Scale bar = 1 mm.

Detailed analysis with two-way ANOVA demonstrated an effect of hemisphere (F_(1,20)_=8.701, p=0.0079) and interaction (F_(1,20)_=11.47, p=0.0029) but not group (F_(1,20)_=2.186, p=0.1548). Bonferroni post-tests showed that there was a significant enlargement of conditioned row B representation in the hM4Di+saline group (### p<0.001) but not in the hM4Di+CNO group (ns p>0.05). Comparison of row B (control or conditioned) width between two groups showed that there was a significant difference in row B width in the conditioned (p<0.01) but not in the control hemispheres (p>0.05) between the two groups. Examples of autoradiograms presenting the whole tangential brain section and magnification of row B representations taken from both tested groups of both hemispheres are shown in **Fig. 5B.**

### L4 SOM-IN inhibition impairs minifreezing to conditioned stimuli

In the course of whisker-based classical conditioning, mice acquire a conditioned response to the conditioned stimulus, which is observed as a decrease in the number of head turns toward CS and was termed “minifreezing”. During the first minute of the first pairing session the mice turn their heads towards the stimulating brush; later this response rapidly decreases (Cybulska-Klosowicz et al., 2009). The percentage of CS accompanied by head movements in response to CS in the hM4Di+saline group was 13.75 ± 3.08% (Day 1), 4.17 ± 1.67% (Day 2), and 5 ± 4.03% (Day 3), and it differed between consecutive days of conditioning (F_(2,17)_=6.671, p=0.0144, repeated measures ANOVA) (**Fig. 6A**, hM4Di+saline). A significant difference was found between Days 1 and 2 (p<0.05) and between Days 1 and 3 (p<0.05) but not between Days 2 and 3 (p>0.05) (Tukey’s multiple comparison tests). These results showed that animals learned that the CS preceded a tail shock. In contrast, analysis of minifreezing in the L4 SOM-IN-inhibited animals showed an impaired behavioral response to the CS with the following percentages of head movements: 62.5% ± 9.83 (Day 1), 52.5% ± 11.44 (Day 2) and 42.5% ± 9.04 (Day 3), and no significant difference in the percentage of head movements in the consecutive days of conditioning was revealed (F_(2,17)_=2.791, p=0.1089, repeated measures ANOVA with Tukey’s multiple comparison tests) (**Fig. 6A**, hM4Di+CNO). To compare the percentage of head movements for all days of the conditioning between both groups, we performed a two-way ANOVA comparison that showed an effect of group (F_(1,30)_=53.64, p<0.0001) but not day (F_(2,30)_=1.916, p=0.1648) or interaction (F_(2,30)_=0.3614, p=0.6997). Bonferroni post-tests showed differences on Day 1 (*** p<0.001), Day 2 (*** p<0.001) and Day 3 (**p<0.01). Altogether, these results indicate that L4 SOM-IN inhibition selectively in the conditioned row of whiskers impairs behavioral response by decreasing minifreezing to conditioned stimuli.

**Fig. 6.**
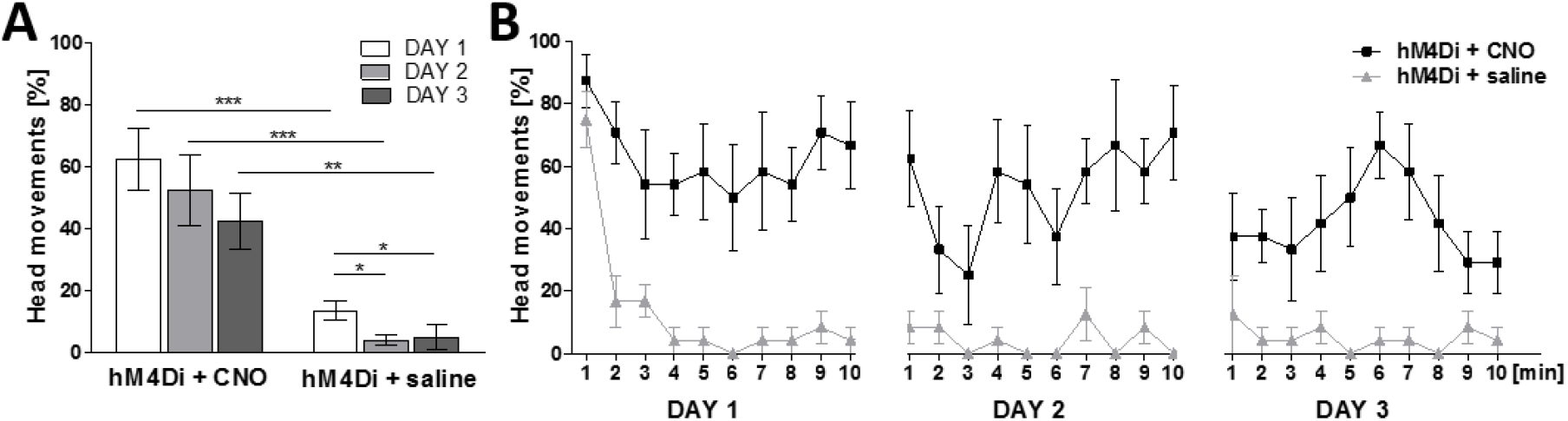
L4 SOM-IN inhibition decreases minifreezing to conditioned stimuli. **A**. Minifreezing shown as a decrease in the percentage of head movements within one day of conditioning (mean±SEM). There was no difference in minifreezing between consecutive days of conditioning in the experimental group (hM4Di+CNO) (F_(2,17)_=2.791, p=0.1089, repeated measures ANOVA with Tukey’s multiple comparison tests, Day 1 *vs* 2: ns p>0.05, 1 *vs* 3: ns p>0.05, 2 *vs* 3: ns p>0.05) but a statistically significant difference in control animals (hM4Di+saline) (F_(2,17)_=6.671, p=0.0144, repeated measures ANOVA with Tukey’s multiple comparison tests, Day 1 *vs* 2: * p<0.05, 1 *vs* 3: * p<0.05, 2 *vs* 3: ns p>0.05). Two-way ANOVA with Bonferroni post-test analysis revealed a significant difference between the control and experimental groups in the percentage of head movements between the corresponding days (F_*(1,30)*_=53.64, p<0.0001): Day 1 (*** p<0.001), Day 2 (*** p<0.001), and Day 3 (*** p<0.01). **B**. Minifreezing shown as a percentage of head movements within every minute of the conditioning in three consecutive days (mean±SEM).

### L2/3 SOM-IN inhibition does not interfere with the formation of plasticity and the behavioral response

L2/3 SOM-INs are important in shaping pyramidal cell activity by targeting their apical dendrites in cortical L1 (Wang et al., 2004). To elucidate the effect of L2/3 SOM-IN activity on the plastic change induced by whisker-based fear conditioning, we selectively blocked L2/3 SOM-INs of the conditioned row representation during conditioning.

Chemogenetic inhibition of L2/3 SOM-INs in row B did not influence the conditioning-induced formation of plasticity. There was a statistically significant increase in the row B representation width between the hemispheres after conditioning (** p=0.0016, two-tailed paired t-test, n=5) **(Fig. 7A)**. The mean width of the conditioned row was 529.5 ± 11.68 μm compared to that of the control row in the opposite hemisphere (438.5 ± 5.64 μm). The plasticity was comparable to that of the control group from previous experiments—DREADD-transduced L4 SOM-INs with saline application (**Fig. 5A, hM4Di+saline**).

The behavioral response to CS application was also examined. The percentage of head movements equaled 20.50% ± 3.20 (Day 1), 9% ± 3.92 (Day 2) and 7.5% ± 2.24 (Day 3). These values were slightly elevated compared to those in the L4 SOM-IN-transduced animals with saline administration (**Fig. 6A, hM4Di+saline**), but no significant differences in the percentage of head movements between days of conditioning within the two groups were revealed (data not shown). To further examine the dynamics of the behavioral response, we analyzed the first three minutes of Day 1 of the conditioning. We found a difference between the first and second as well as between the first and third but not between the second and third minute of the conditioning (F_(2,14)_=25.57, p=0.0003, repeated measures ANOVA with Tukey’s multiple comparison tests; minute 1 *vs* 2: ** p<0.01, 1 *vs* 3: *** p<0.001 and 2 *vs* 3: ns p>0.05) (**Fig. 7B**). These results suggest that acquiring the behavioral response occurs at the very beginning of the conditioning and that minifreezing is stable across all sessions.

**Fig. 7.**
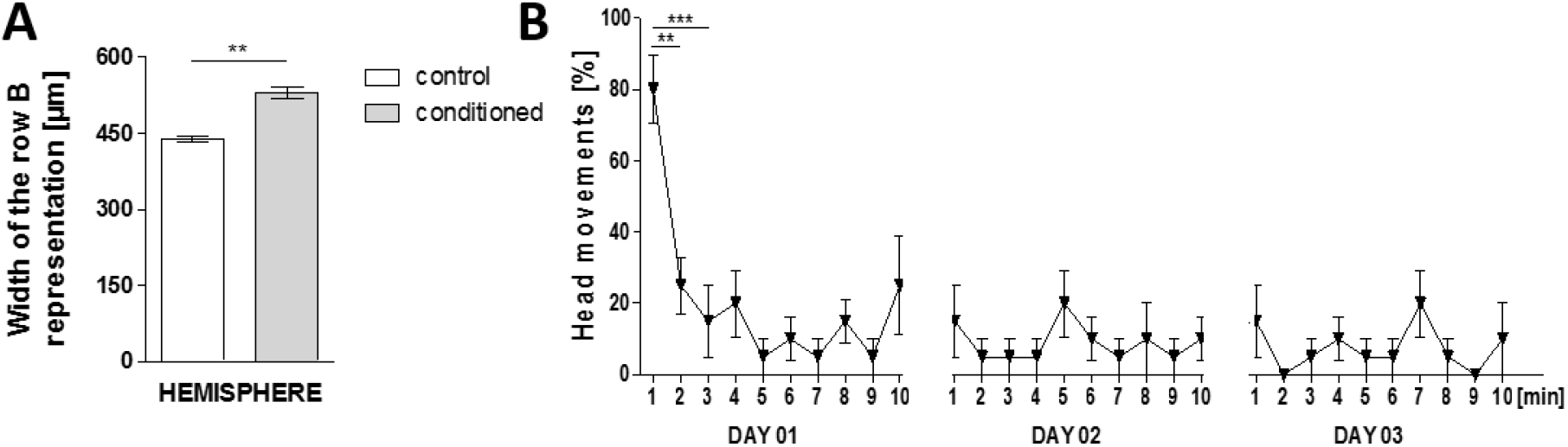
L2/3 SOM-IN inhibition does not interfere with the formation of plasticity and the behavioral response. **A**. Row B representation width [μm] of the control and conditioned hemispheres in animals with L2/3 SOM-IN inhibition during conditioning (mean±SEM). Two-tailed paired t-tests showed a significant increase in row B width after conditioning (** p=0.0016, n=5). **B**. Minifreezing showed as a percentage of the head movements within every minute of the conditioning in three consecutive days (mean±SEM). Repeated measures ANOVA with Tukey’s multiple comparison tests revealed significant differences in the number of head movements between the first three minutes of Day 1 (F_(2,14)_=25.57, p=0.0003). Minute 1 *vs* 2: ** p<0.01, 1 *vs* 3: *** p<0.001 and 2 *vs* 3: ns p>0.05.

Together, these results indicate that L2/3 SOM-INs do not affect the formation of plasticity and do not impair the behavioral response to CS during whisker-based classical conditioning.

### L4 SOM-IN chemogenetic excitation does not influence learning-induced plasticity

Increasing L4 SOM-IN activity may decrease the activity of PV-INs, resulting in increased activity of excitatory neurons (Xu et al., 2013), and increased SOM-IN activity during learning has been reported earlier (Cummings and Clem, 2020; Cybulska-Klosowicz et al., 2013b). Moreover, SOM-IN transplantation has been shown to drive cortical plasticity and reshape neuronal networks (Tang et al., 2014). We aimed to determine how chemogenetic activation of L4 SOM-INs during conditioning affects plasticity induced by whisker-based fear learning.

Chemogenetic stimulation of L4 SOM-INs in row B did not influence the conditioning-induced plasticity since in both groups (hM3Dq+CNO, n=5, and hM3Dq+saline, n=4), plasticity was induced to a comparable level (1.182 ± 0.029 in hM3Dq+CNO and 1.193 ± 0.03 in hM3Dq+CNO group; ns p=0.9048 two-tailed Mann-Whitney test) (**Fig. 8A**). Two-way ANOVA with Bonferroni post-tests revealed an effect of hemisphere (F_(1,14)_=30.78, p<0.0001) but not group (F_(1,14)_=0.5612, p=0.4662) or interaction (F_(1,14)_=0.01804, p=0.8951), confirming a significant difference in the width of row B labeling between the two hemispheres in both the hM3Dq+CNO (## p<0.01) and hM3Dq+saline (## p<0.01) groups. We observed consistent labeling of the row B representation in both hemispheres: conditioned (p>0.05) and control (p>0.05). Example autoradiograms presenting the whole tangential brain section and magnification of row B representations taken from both tested groups of both hemispheres are shown in **Fig. 8B**. Since the learning of stereotyped movements was interrupted after not only inhibition but also excitation of SOM-INs (S. X. Chen et al., 2015), we aimed to determine the behavioral response in the hM3Dq+CNO mice. L4 SOM-IN activation did not influence the conditioned response. Two-way ANOVA revealed an effect of day (F_(2,18)_=4.304, p=0.0297) but no effect of group (F_(1,18)_=0.03297, p=0.8580) or interaction (F_(2,18)_=0.03915, p=0.9617). Bonferroni post-tests showed no difference in the number of head movements between the hM3Dq+CNO and hM3Dq+saline groups in the course of conditioning: Day 1 (ns p>0.05), Day 2 (ns p>0.05) and Day 3 (ns p>0.05). (**Fig. 8C**). Since no difference between the two groups in the conditioned response was observed, behavioral data of the hM3Dq+CNO group indirectly confirmed that CNO (1 mg/kg) administration does not influence behavioral response *per se*, thus validating the results from the SOM-IN inhibition experiment. Together, these behavioral results are in line with the 2DG data, indicating no effect of L4 SOM-IN excitation on whisker-based fear learning.

**Fig. 8.**
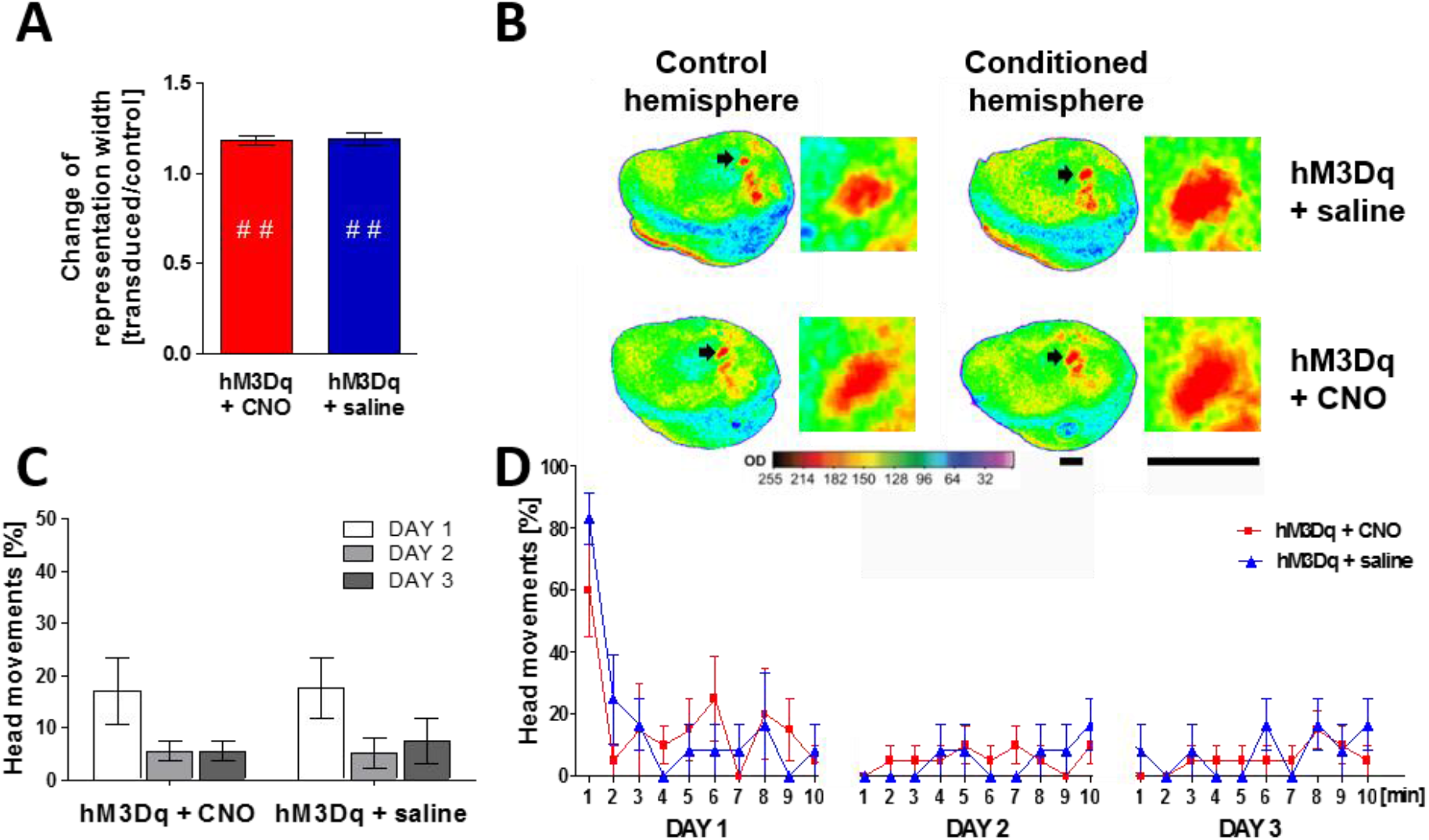
L4 SOM-IN chemogenetic excitation does not influence learning-induced plastic change formation or minifreezing to conditioned stimuli. **A.** Plasticity shown as a change in representation width, calculated as a ratio between a representation width of conditioned (transduced) row B to a control one in the opposite hemisphere (mean±SEM). Two-way ANOVA with Bonferroni post-test analysis showed a significant difference in row B width between the control and conditioned hemispheres (F_(1,14)_=30.78, p<0.0001) in the control group (hM4Di+saline) (## p<0.01, n=4) and in the experimental group (hM3Dq+CNO) (## p<0.01, n=5). No significant difference in plasticity between the two groups (ns p=0.9048, two-tailed Mann-Whitney test) was observed. **B.** Examples of representative, pseudocolored autoradiograms (a pair of left control and right conditioned hemisphere shown in a row) taken from both the control (hM3Dq+saline) and experimental groups (hM3Dq+CNO). Black arrows indicate row B representations magnified in a square on the right side. Scale bar = 1 mm. **C.** Minifreezing is shown as a percentage of head movements within one day of conditioning (mean±SEM). Two-way ANOVA with Bonferroni post-test analysis revealed no difference between the control and experimental groups in the percentage of head movements between the corresponding days (F_(1,18)_=0.03297, p=0.8580): Day 1 (ns p>0.05), Day 2 (ns p>0.05), and Day 3 (ns p>0.05). **D.** Minifreezing is shown as a percentage of the head movements as a function of time (mean±SEM).

### L2/3 VIP-IN inhibition does not affect the L4 SOM-IN-mediated plasticity

Recent studies have shown that interneurons containing vasoactive intestinal polypeptide play a key role in a disinhibitory circuit that regulates adult visual cortical plasticity (Fu et al., 2015) or associative learning in the auditory cortex (Pi et al., 2013) by blocking somatostatin-containing interneurons (VIP-SOM circuit). L2/3 VIP-INs are highly active during whisking (Lee et al., 2013), and since they preferentially innervate SOM-INs, we asked whether they can play a role in shaping L4 SOM-IN-related plasticity. We found that blocking L2/3 VIP-INs in row B cortical representation during whisker-based associative learning did not influence the induction of plasticity, and its extent was similar in both tested groups (1.19 ± 0.01 in hM4Di+CNO and 1.19 ± 0.01 in hM4Di+saline; ns, p=0.9524, two-tailed paired t-test, n=6 per group) (**Fig. 9A**). Two-way ANOVA with Bonferroni post-tests of row B labeling width revealed a main effect of hemisphere (F_(1,20)_=70.97, p<0.0001) and group (F_(1,20)_=7.847, p=0.0110) but not interaction (F_(1,20)_=0.05957, p=0.8097). Post-tests confirmed the interhemispheric difference in row B widths in both analyzed groups (p<0.001) with no significant difference in row B labeling width between the groups (ns p>0.05 for both hemispheres). Examples of autoradiograms presenting the whole tangential brain section and magnification of row B representations taken from both hemispheres of tested groups are shown in **Fig. 9B**. We also analyzed the conditioned response to CS in both groups, and the results showed no effect of L2/3 VIP-IN inhibition on the behavioral response; the data were comparable to the results obtained in the SOM-IN excitation experiment (**Fig. 8C** and **D**) (data not shown). In conclusion, the obtained data suggest no effect of L2/3 VIP-IN activity on the formation of plastic change induced by whisker-based classical conditioning.

**Fig. 9.**
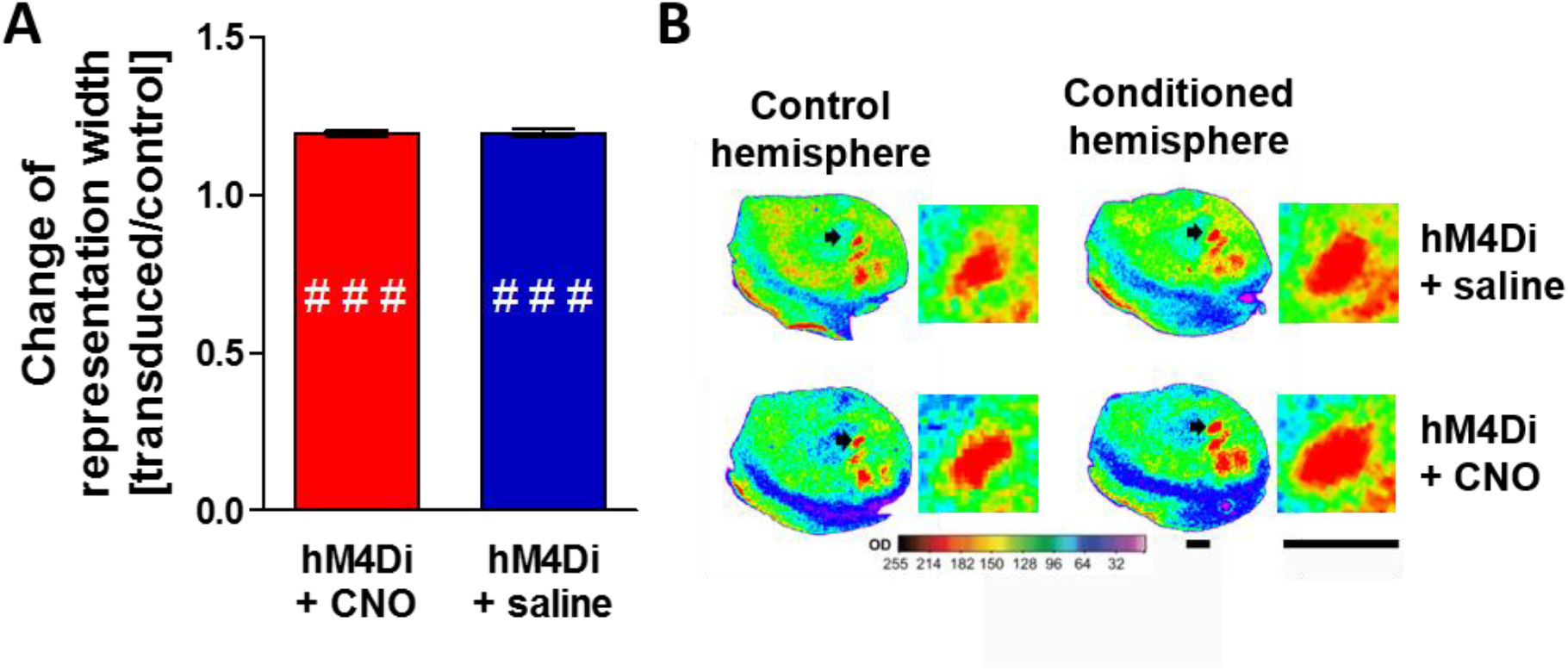
L2/3 VIP-IN chemogenetic inhibition does not influence the whisker-based learning-induced plastic change. **A.** Plasticity shown as a change in representation width, calculated as a ratio between a representation width of conditioned (transduced) row B to a control one in the opposite hemisphere (mean±SEM). Two-way ANOVA with Bonferroni post-tests showed a significant difference in row B width between the control and conditioned hemispheres (F_(1,20)_=70.97, p<0.0001) in the control group (hM4Di+saline) (### p<0.001, n=6) and in the experimental group (hM4Di+CNO) (### p<0.001, n=6). No significant difference in plasticity between the two groups (ns p=0.9524, two-tailed unpaired t-test) was observed. **B.** Examples of representative, pseudocolored autoradiograms (a pair of left control and right conditioned hemispheres shown in a row) taken from both the control (hM4Di+saline) and experimental groups (hM4Di+CNO). Black arrows indicate row B representations magnified in a square on the right side. Scale bar = 1 mm.

## DISCUSSION

We found that chemogenetic inactivation of L4 SOM-INs during whisker-based classical conditioning impaired the learning and plasticity of cortical representation. This result was layer-specific, as silencing L2/3 SOM-INs did not affect learning or learning-induced plasticity.

We previously found indications that SOM-INs are involved in learning-dependent cortical reorganization (Cybulska-Klosowicz et al., 2013b). Here, we showed that they are necessary for the development of plastic change in the cortex and normal development of the conditioned reaction. Numerous experiments implementing different fear conditioning paradigms have shown the involvement of cortical processing in learning (Banerjee et al., 2017; Gillet et al., 2018; Letzkus et al., 2011; Weinberger, 2015). In the barrel cortex, conditioning resulted in changes in strength in intercolumnar and intracolumnar connections (Rosselet et al., 2011; Urban-Ciecko et al., 2005). It is, however, known that the most likely location of learning-induced plastic modifications are supragranular cortical layers; thus, it would be reasonable that alterations in L2/3 circuits would influence cortical reorganization, while in our experimental paradigm, normal functioning of L4 but not L2/3 SOM inhibitory circuits was critical. Therefore, it seems that the outcome we observed would be a specific result of the local impact of L4 SOM-INs on the L4 neuronal network, affecting the ability of cortical reorganization as a consequence of fear conditioning. Neurons in somatosensory L4 constitute a unique network, with the majority of excitatory cells in S1 being stellate cells and SOM-INs being non-Martinotti cells integrated into the local circuit within L4 (Scala et al., 2019). In terms of connectivity, L2/3 and L4 SOM-INs also differ. In L2/3, SOM-INs inhibit mainly pyramidal cells, whereas in L4, they preferentially suppress PV-INs. L4 SOM-INs can decrease the feedforward inhibition of excitatory neurons by fast-spiking PV-INs and disinhibit the transmission of the afferent signal from the thalamus (Xu et al. 2013). We hypothesized that in response to the conditioned sensory stimulus (whisker activation) and UCS, this disinhibitory effect can be augmented, more effectively removing feed-forward inhibition of excitatory neurons during sensory input (Liguz-Lecznar et al., 2016). In this way, the mechanism of CS+UCS upon L4 circuitry would rely on removing the gating of the thalamocortical signal by PV interneurons. UCS-driven cholinergic projection to the cortex acts on PV cell synapses onto principal neurons *via* inhibitory M2 cholinergic receptors, effectively weakening the inhibition of principal cells by PV neurons (Kruglikov and Rudy, 2008). Cholinergic input to SOM-INs, *via* both muscarinic and nicotinic receptors, is effective at much lower agonist doses than those in other interneurons (N. Chen et al., 2015). Activation of SOM-INs by acetylcholine may contribute to inhibition of PV interneurons and consequently to stronger disinhibition of principal cells. Moreover, touch-induced activation of SOM-INs, including L4, is time delayed, probably reflecting intracortical excitatory input rather than thalamic input (Yu et al., 2019); thus, L4 SOM-INs may increase their spike rates as a result of increased activity of excitatory cells during conditioning, further potentiating L4 disinhibition. SOM-PV disinhibition in the thalamocortical input layer during CS+UCS pairing may allow for wider spreading of the signal from the active vibrissae in the barrel cortex. A similar disinhibitory circuit was proposed to underlie social fear in the dorsomedial prefrontal cortex, and it was shown that SOM-IN inhibition alleviates the behavioral social fear response in conditioned mice and reduces the firing rates of PV-INs, which are elevated during social fear expression (Xu et al., 2019).

In our experiments, the inhibition of L2/3 SOM-INs did not interfere with plasticity induction. The effect of conditioning, as seen with 2DG, is primarily in L4; the changes in neuronal interactions that were observed in L2/3 (Lebida and Mozrzymas, 2017; Rosselet et al., 2011; Urban-Ciecko et al., 2005) are not reflected in the expansion of the trained whisker representation in this layer. Moreover, in the somatosensory cortex, Denardo et al. (2015) found strong input from L3 mainly to L6; therefore, the direct and powerful effect of L2/3 SOM-IN activity alteration on L4 is rather unlikely. However, Xu et al. (2013) have shown that in the sensory cortex, optogenetic inhibition of SOM-INs in an active cortical network increased the firing of L2/3 principal cells, and here, we expected a similar effect but with chemogenetics. Thus, the effect of L2/3 SOM-IN inhibition on the ability of L4 functional reorganization, if any, could be supportive rather than suppressive for induction of plastic change, targeting supragranular layers directly *via* the local L2/3 network instead of acting via disinhibition of the L4 afferent signal.

A previous reports showed that 2DG labeling is largely ascribed to synaptic activity (Schwartz et al., 1979), so the signal in L4 may originate from thalamocortical axons and interbarrel synaptic connections (Lübke et al., 2003; Meyer et al., 2010). Thus, alternatively, we could interpret the result observed in L4 as a manifestation of the plasticity of the thalamocortical input itself, which has been demonstrated in paradigms not involving associative learning (Chung et al., 2017; Oberlaender et al., 2012; Wimmer et al., 2010) or as a passive reflection of learning-dependent plasticity at the subcortical level. We have no data about learning-dependent plasticity in the barreloids, but in a conditioning paradigm involving all whisker stimulation, we found that learning increased correlations between structures of the thalamocortical loop and increased activity of both thalamic ventral posterior and posterior medial nucleus (Cybulska-Klosowicz et al., 2013a). Nevertheless, the possibility that what we observed in cortical L4 is solely the reflection of subcortical changes seems implausible since our experimental paradigm induced many electrophysiological, biochemical, and molecular changes in the cortex. Previously, we found, in barrels representing the trained whiskers in L4, inhibitory synaptogenesis and morphological indices of local protein synthesis in spines, long-lasting alterations in NMDA receptor subunit composition, GAD mRNA and protein expression, and changes in neuronal network activity as well as in intrinsic excitability and tonic and phasic GABA currents (Jasinska et al., 2016; for review see Liguz-Lecznar et al., 2016; Siucinska, 2019).

Interestingly, contrary to a study in which both suppression and enhancement of SOM-IN activity in the motor cortex altered synaptic plasticity and impaired learning (S. X. Chen et al., 2015), we found that enhanced excitation of L4 SOM-INs failed to modify fear learning-induced plasticity. It is plausible that during conditioning, the level of L4 SOM-IN activity in cognate barrels is already high enough to enable principal neuron disinhibition and spread the signal across the column, and further excitation does not make a difference. Recently, excitatory DREADD transduction was shown to change basic activity without CNO administration, probably *via* leaky signaling that led to chronic low-level activation inducing compensatory effects (Rosenthal et al., 2020); however, we did not observe any altered 2DG labeling in any group of hM3Dq-transduced animals.

L2/3 VIP-IN chemogenetic inhibition during conditioning also failed to modulate learning-induced plastic changes in L4. It was shown that with passive whisker deflection, touch increased the spike rate of L2/3 VIP-INs, which disinhibited excitatory neurons and PV-INs in the barrel cortex, likely *via* inhibition of SOM-INs (Yu et al., 2019). The inhibition of L2/3 VIP-INs should therefore disinhibit SOM-INs; thus, we conclude that in our experimental paradigm, neither direct inhibition nor indirect disinhibition of L2/3 SOM-INs affect L4 plasticity. The most plausible explanation of the latter effect involves connectivity of VIP-INs: although the axonal tree of L2/3 VIP-INs is observed across all layers of the mouse barrel cortex, including L4, their axonal boutons are predominant in L2/3 and 5a, being rather rare in L4, at 42.8%, 22.3% and 13.3% (Prönneke et al., 2015). Additionally, in L4, only 34% of the VIP-IN axonal boutons are located on GABA+ dendrites, suggesting weak inhibitory-inhibitory interactions in this layer (Zhou et al., 2017). Muñoz et al. (2017) have shown stronger synaptic inputs from VIP-INs onto L2/3 SOM-INs than onto L4 to L6 SOM-INs. To the best of our knowledge, there is no direct evidence of the functional existence of the VIP to L4 SOM circuit, so VIP-IN activation fails to modulate L4 SOM-IN-dependent plasticity. Data concerning the involvement of VIP-IN activity in learning and plasticity are contradictory. On the one hand, VIP-IN activation enhances adult visual cortical plasticity (Fu et al., 2015), and whisker-based object localization in the go/no-go task increases the spike rates of VIP-INs in the barrel cortex (Yu et al., 2019). On the other hand, the selectivity of VIP-INs in the visual go/no-go discrimination task did not change in the course of associative learning (Khan et al., 2018).

Taken together, our study revealed that, in contrast to L2/3 SOM-INs activity, the activity of L4 SOM-INs during conditioning was indispensable to produce learning-induced plasticity in the barrel cortex after classical conditioning involving whiskers. Both L4 SOM-IN excitation and L2/3 VIP-IN inhibition did not interfere with the induction of plasticity. We hypothesize that L4 SOM-INs may act as regulators of the PV-INs’ gating of thalamic input and, upon neuromodulatory input, may enhance their activity to increase local excitation, allowing the flow of information across the cortical column.

Our results indicate that interactions between thalamic input and L4 neuronal circuits are the basis for initiation of the effect of learning-evoked cortical plasticity of whisker cortical representation. The activity of SOM interneurons appears to be a critical element of these L4 circuits.

## Acknowledgments

Experiments were supported by a Polish National Science Centre Grant to MK (2015/17/B/NZ4/02016).

## Notes

Conflict of interest statement: The authors declare no financial and non-financial competing interests.

### Competing Interest Statement

The authors have declared no competing interest.

